# Low-frequency Cortical Activity Reflects Context-dependent Parsing of Word Sequences

**DOI:** 10.1101/2024.11.12.623335

**Authors:** Honghua Chen, Tianyi Ye, Minhui Zhang, Nai Ding

**Author notes:** **Corresponding author:** Nai Ding. **Author contributions:** Conceptualization, H.C., T.Y., and N.D.; formal analysis, investigation, and project administration, H.C., M.Z., and N.D.; writing - original draft, H.C.; writing – review and editing, all authors.

## Abstract

During speech listening, it has been hypothesized that the brain builds representations of large linguistic structures such as sentences, which are captured by neural activity tracking the rhythm of these structures. Nevertheless, it has been concerned that the brain may only encode words, and neural activity tracking structures may be confounded by neural activity tracking the predictability or syntactic properties of individual words. Here, to disentangle the neural responses to sentences and words, we design word sequences that are parsed into different sentences in different contexts. By analyzing neural activity recorded by magnetoencephalography, we find that low-frequency neural activity strongly depends on the context – The difference between MEG responses to the same word sequence in two contexts yields a low-frequency signal, most strongly generated in the superior temporal gyrus, which precisely tracks sentences. The predictability and syntactic properties of words can partly explain the neural response in each context but cannot explain the difference between contexts. In summary, low-frequency neural activity encodes sentences and can reliably reflect how the same word sequence is parsed in different contexts.

## Introduction

Human language organizes words into phrases and sentences (Chomsky, 1965; Everaert et al., 2015), and complex neural computations are required to integrate words and derive the compositional meaning of a sentence (Hagoort, 2019; Pylkkänen, 2019). It has been proposed that neural activity on different time scales separately encode linguistic units of various sizes, including syllables (Mesgarani et al., 2014), words (Ding et al., 2018), sentences (Ding et al., 2016; Woolnough et al., 2023), and discourses (Lerner et al., 2011). More specifically, neural activity encoding a linguistic unit is hypothesized to track the unit – Neural activity can signify the onset and offset of the unit and may keep track of how long the unit has lasted at each moment (Ding, 2020; Lakatos et al., 2019). Critically, it has been suggested that neural encoding of larger linguistic structures such as phrases and sentences is not confounded by neural encoding of prosodic cues associated with these structures, since the phrase- or sentence-tracking activity remains when the prosodic cues are deprived (Ding et al., 2016).

Nevertheless, it has been concerned that sentence-tracking activity may still be confounded by word-level features, e.g., the predictability of individual words (Goldstein et al., 2022; Heilbron et al., 2022; Weissbart & Martin, 2024) or the syntactic properties of individual words (Frank & Yang, 2018; Kalenkovich et al., 2022) – Within a sentence, initial words are less predictable than later words. Furthermore, a sentence usually begins with a noun phrase and in many experiments investigating sentence-tracking activity, the sentences always start with a noun phrase and have uniform syntactic structures. Therefore, neural activity encoding word predictability or syntactic properties may appear to track sentences. To tease apart neural activity encoding words and sentences, several studies design stimuli in which words alternating in their part-of-speech information construct sentences or not (Burroughs et al., 2021; Lo et al., 2022; Lu et al., 2022, 2023). For instance, alternations between adjectives and nouns construct phrases (e.g., loud room) while alternations between adjectives and verbs do not construct phrases (e.g., rough tell), and it is found that only the adjective-noun condition generates a phrase-tracking response (Burroughs et al., 2021). A concern for these studies is that words presented in a phrasal context may receive deeper processing (Gwilliams et al., 2023; Woolnough et al., 2023) than words that do not group into phrases.

Here, to disentangle the neural tracking of sentences and words, we designed ambiguous word sequences that can be grouped into different sentences in different conditions (Fig. 1a). The ambiguous word sequences are designed based on the relatively flexible word order in Chinese. In the two conditions, termed the subject-verb-object (SVO) and the object-subject-verb (OSV) conditions, the words are separately grouped into SVO and OSV sentences, which have distinct sentence boundaries. Native Chinese speakers listened to the word sequences while their neural responses were recorded using magnetoencephalography (MEG). We analyze the MEG response to the word sequence that is shared by the SVO and OSV conditions (shaded area in Fig. 1). Since the word sequence is identical in the SVO and OSV conditions, neural activity encoding syntactic properties of individual words should also be identical in both conditions and get removed in the difference response between the two conditions (Fig. 1c). The only word property that may differ between the two conditions is word predictability since the OSV condition has an additional word at the beginning. In the following, we will use the GPT-2 model to investigate whether word predictability differs between conditions. If not, neural activity encoding word predictability is also removed in the difference response. Neural activity tracking sentence onsets, however, is preserved in the difference response since the sentence onsets differ between conditions.

**Fig. 1.**
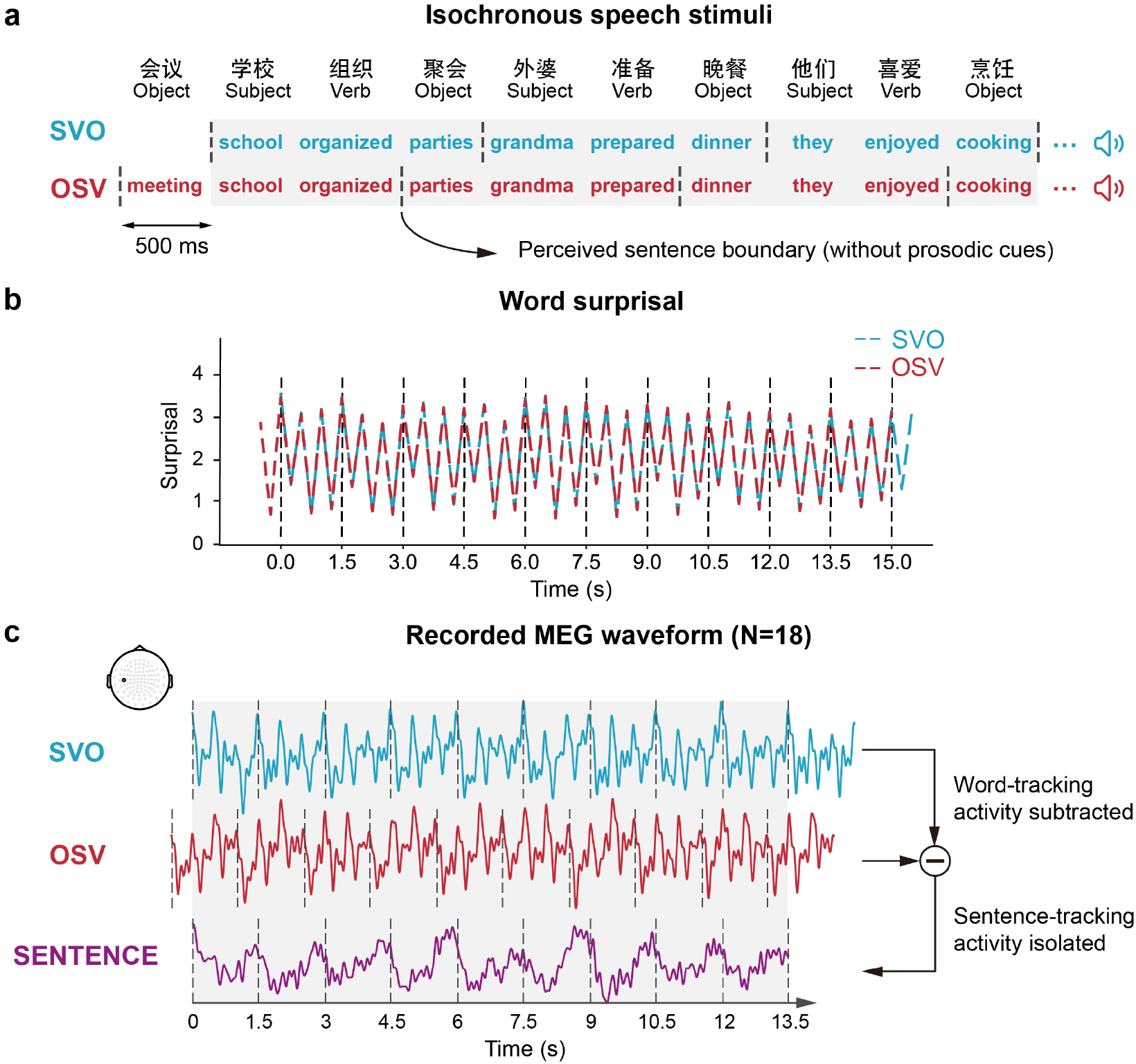
Experimental design and analysis method. **a** Participants listen to sentences with two distinct word orders, i.e., SVO and OSV. In SVO condition, the words organize into 12 SVO sentences. The OSV condition is the same as the SVO condition except that an additional word is added to the beginning while the last word is removed. Each word contains 2 monosyllabic morphemes and lasts 500 ms so that the sentences and words are presented in 2/3 Hz and 2Hz respectively. In all analyses, the first word in the OSV condition and the last word in the SVO condition are removed, so that the two conditions share exactly the same word sequence, which is shaded in gray. **b** The surprisal of each morpheme is calculated using the GPT2 model. The surprisal sequence is highly consistent between conditions during the common word sequence (shaded in gray). **c** MEG response grand averaged over 18 participants. We select a representative sensor in the left temporal region and the response is filtered below 10 Hz. For the neural responses to the common word sequence (shaded in gray), the difference response between SVO and OSV conditions is referred to as the SENTENCE response.

By analyzing the difference response between SVO and OSV conditions, the following analyses demonstrate that low-frequency neural activity faithfully tracks sentences and reflects context-dependent parsing of word sequences. Furthermore, the strength of sentence-tracking activity is correlated with the performance of a sentence comprehension task. A representational similarity analysis (RSA) further reveals that, within the SVO or OSV condition, neural neural activity encodes sentences (i.e., ordinal position of a word within a sentence), syntactic properties of words (i.e., part of speech, PoS), and predictability of words (i.e., word surprisal) with different temporal patterns.

## Results

### Context-dependent Neural Parsing of Word Sequences

We first illustrated the trial-average MEG waveform using a representative MEG sensor which is located in the left temporal region (Fig. 1c). The responses in the SVO and OSV conditions both exhibited responses evoked by individual words, i.e., three response peaks within the dashed lines illustrating sentence boundaries, and responses tracking sentences, i.e., the consistent slow drift within the dashed lines. When the response waveform was subtracted between the SVO and OSV conditions, which is referred to as the SENTENCE response, the responses to individual words were effectively cancelled and what remained was a slow response that repeated every ∼1.5 s, i.e., the duration of a sentence. A concern for this isolation is the additional word in OSV at the beginning, which might alter the word predictability. We therefore use the GPT-2 model to calculate the predictability and find the surprisal, i.e., the negative logarithm of probability, is highly consistent between SVO and OSV (Fig, 1b) and has no effect at the sentence rate. Thus, the SENTENCE response is hypothesized to contain only the neural response to sentences.

Next, we characterize the MEG response in the frequency domain. Since the power spectrum of trial-averaged, i.e., evoked, MEG response was always positive and was not influenced by the polarity of the response, we averaged the response power spectrum across all gradiometers. The averaged spectrum showed clear peaks at both sentential and word rates in both SVO and OSV conditions (Fig. 2a). The response power at the sentential and word rates was significantly stronger than the power averaged over neighboring frequencies in both conditions (*P* < 0.001 for all tests, two-sample paired *t*-test, FDR corrected). For the difference waveform between SVO and OSV conditions, the power spectrum showed a significant peak at the sentence rate (Fig. 2a, *P* = 0.0021, two-sample paired *t*-test, FDR corrected) but not at the word rate (Fig. 2a, *P* = 0.60, two-sample paired *t*-test).

**Fig. 2.**
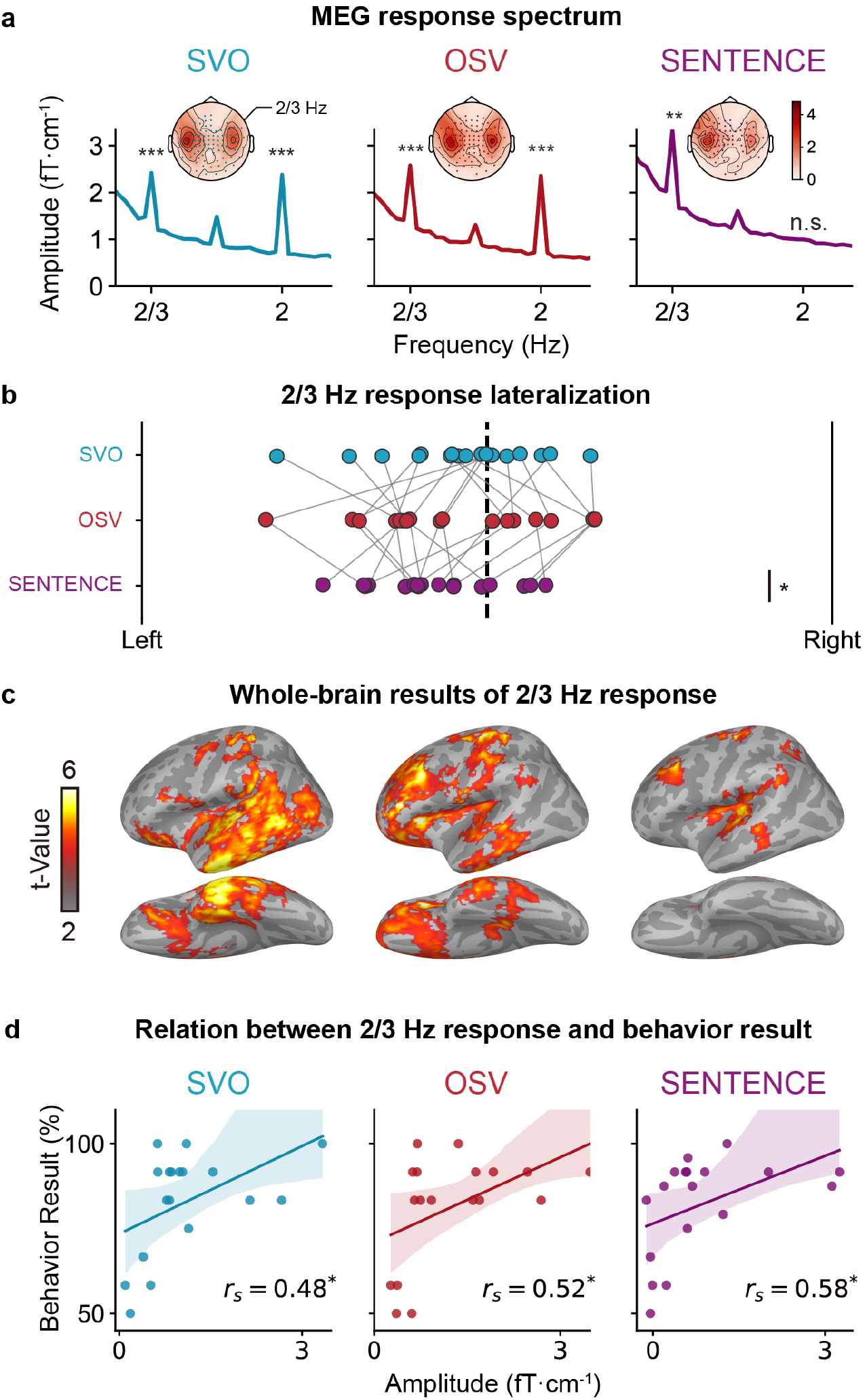
Low-frequency neural response in the four conditions. **a** Lines show the MEG response spectrum averaged over participants and sensors. Frequency bins with significantly stronger amplitude than estimated noises are marked (** *P* < 0.01, *** *P* < 0.001, one-sided t-test, FDR corrected). Topographies show maps of 2/3 Hz response amplitude across sensors. Dotted sensors show significant responses (threshold at permutation t-test *P* < 0.01 across 5000 permutations, FDR corrected). **b** The lateralization index (LI = R /(L+R)) of the 2/3 Hz response across participants (* *P* < 0.05, two-sided t-test). **c** The whole-brain source-level activity of 2/3Hz response amplitude (Threshold at *P* < 0.05 using a cluster-based permutation test, with cluster forming using threshold-free cluster enhancement (TFCE) and the number of permutations = 10000). **d** The relation between normalized 2/3 Hz response and behavior results across participants. The relation was evaluated via Spearman correlation. * *P* < 0.05.

### Neural Sources of Sentential-rate Responses

For both SVO and OSV conditions, as well as the difference waveform, the sensor-space MEG topography of sentential-rate response showed bilateral activations (Fig. 2a), with the difference waveform significantly left-lateralized (Fig. 2b, lateralization index = 0.42, P = 0.0069, one-sample t-test, FDR corrected). We further localized the neural sources of the sentential-rate responses in the left hemisphere (Fig. 2c, threshold at *P* < 0.05, TFCE corrected with 10,000 nonparametric permutations). In SVO and OSV, the sentential-rate response showed widespread activation across the canonical language network, including left frontal and temporal cortices. The contrast between SVO and OSV showed no significant difference. The difference waveform showed activations in the middle superior temporal gyrus (STG), the posterior temporal cortex (PTC) and the middle frontal gyrus (MFG).

### Correlation between Sentential-rate Activity and Behavior

We further tested the relationship between MEG responses and behavioral results (Fig. 2d). In the experiment, participants were asked to detect occasionally presented semantically abnormal sentences (e.g., “the newspaper breaks the sky”) after each trial. The frequency of successfully reporting abnormal sentences in outlier trials was 82.9% ± 15.3% and 81.5% ± 16.1% (mean ± std.) in the SVO and OSV conditions respectively (no significant difference between conditions, *P* = 0.61, two-sample paired *t*-test). The sentential-rate responses in the SVO and OSV conditions, as well as in the difference waveform, significantly correlated with the behavioral result (SVO: Spearman’s R = 0.48, P = 0.039; OSV: Spearman’s R = 0.52, P = 0.039; SENTENCE: Spearman’s R = 0.58, P = 0.033, FDR corrected).

### Neural Encoding of Word Property and Sentence

The previous analyses focused on the sentence-tracking response, but in the SVO and OSV conditions a word response was also clearly visible (Fig. 1c). Next, we investigated the temporal dynamics of neural activity that separately encoded word properties and sentences. We focused on two word properties, i.e., PoS and word surprisal, and the sentence property was the ordinal position of a word within the sentence, which was 1, 2, or 3 in the current experiment. We characterized how the three speech features, i.e., PoS, surprisal, and word order, was encoded in the multichannel MEG activity using the representational similarity analysis (RSA) (Kriegeskorte et al., 2008; Lyu et al., 2019). We calculated the representational dissimilarity matrices (RDMs) among 120 words (4 averaged trials and 30 words per trial) we presented during the experiment, for both MEG responses and the three speech features (Fig. 3a). We used partial Pearson correlation to assess the fit level of these model RDMs for neural RDMs. In addition, we controlled the neural representation of phonetics and conditions using two control RDMs (Fig. S2a).

**Fig. 3.**
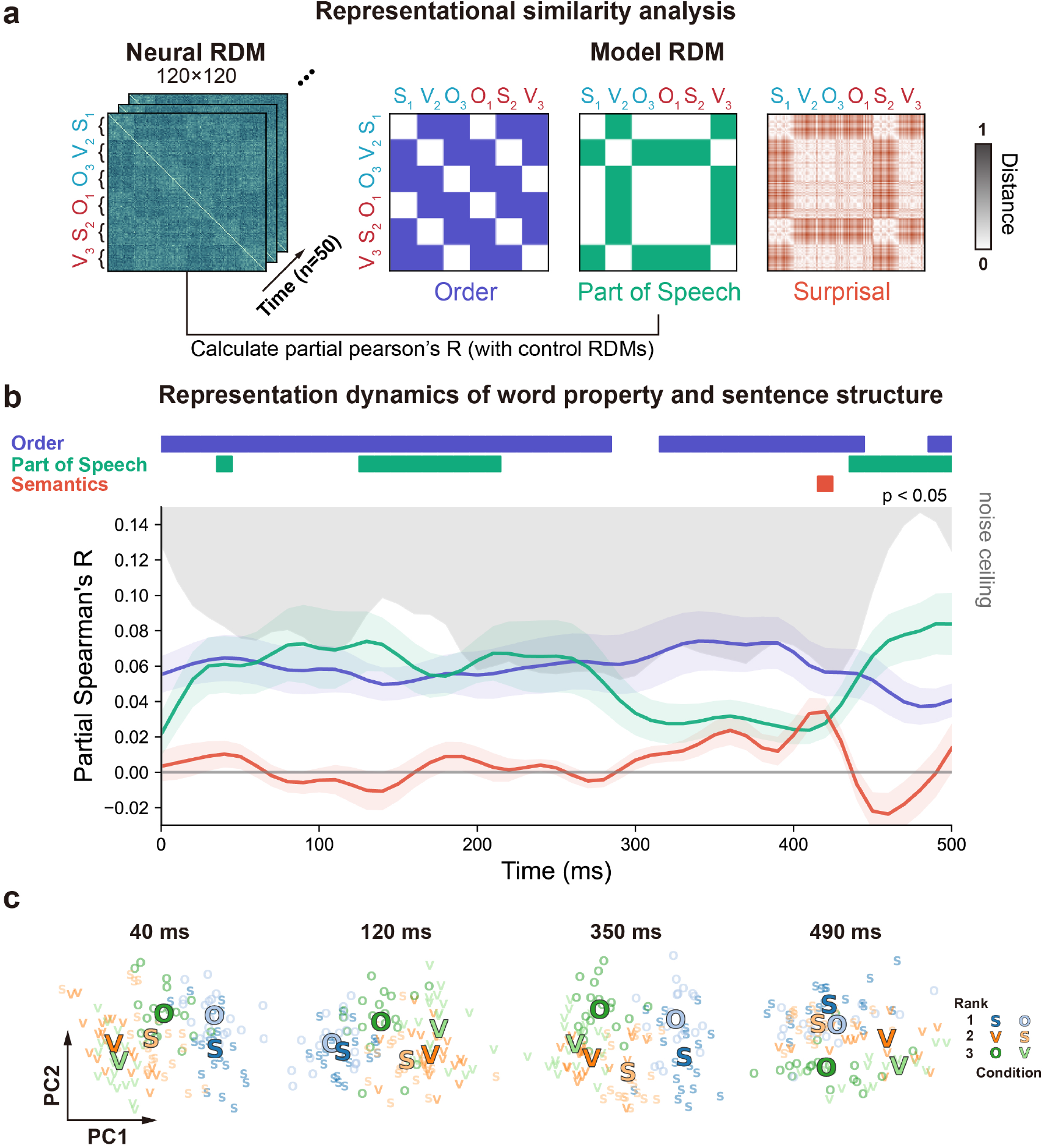
Representational similarity analysis (RSA) revealed the temporal dynamics of word property and sentence structure representation. **a** Illustration of the pipeline for time-resolved RSA. The neural representational dissimilarity matrices (RDMs) were calculated by deriving the neural responses of each word (averaged over trials) and computing the pair-wise correlation. As a word lasted 500 ms, we calculated the RDM at each time window of 10 ms and got 50 neural RDMs. We then correlated the neural RDMs with three hypothesized model RDMs. Considering the correlation between model RDMs (Fig. S2b), we used the partial Pearson correlation here. We further added two control RDMs to control the effects of condition-specific and phonetic processing (Fig. S2a). **b** Time-resolved model fit. The line chart shows the partial Pearson correlation between whole-brain neural RDMs and three model RDMs. The shaded area covers 2 SEM over participants (*N*=18). Significant time points are labeled above the plot (threshold at *P* < 0.05 using a cluster-based permutation test, with cluster forming at point-wise t-test *P* < 0.001 and number of permutations = 5000). The grey shaded area shows the noise ceiling. The model fit for control RDMs is shown in Fig. S2c. **c** PCA plots of the RDM data, at four representative time points. Each point shows an item in the RDM, and the rimmed points show the centers of the 6 classes of items. The rank of each item in the sentence is represented by 3 hues. Items in dark and light colors are in SVO and OSV respectively.

The RSA analysis was applied to each time point in a 500-ms time window starting from the onset of a word. Figure 4b shows the time-resolved model fit throughout the word duration. The neural response was significantly explained by the word order within a sentence across the whole 500 ms (threshold at *P* < 0.05, corrected using a cluster-based permutation test with cluster forming at point-wise *t*-test). The PoS of words significantly explained the neural response at 120∼250 ms and 450∼500 ms after the word onset. The surprisal model significantly explained the neural response at around 420 ms after the word onset. Besides, the neural responses representing the phonetics showed a peak at around 140 ms (Fig. S2c).

We use principal component analysis (PCA) to visualize the RDMs at these time points (Fig. 3c). Closer points show similar dissimilarity patterns of the RDMs. The plots are consistent with the RSA results. Nouns (S and O in the plots) and verbs were clustered at 120 ms and 490 ms, while words of different ranks are clustered around the entire period and best separated at 350 ms. Overall, these results reveal the temporal dynamics of linguistic representation underlying the low-frequency neural responses.

## DISCUSSION

Sentence understanding needs intricate computation to access the words and organize them into sentences, yet the underlying neural implementation remains unsettled. It has been proposed that neural activity at the time scale of sentence tracks the sentence structure. However, the sentence-tracking activity is confounded by the neural responses to the properties of individual words including the syntactic feature and the lexical predictability. The current study proposes a new method to isolate the sentence-level neural response by comparing two conditions with the exactly same word sequences but different sentence boundaries. The results demonstrate the content-dependent parsing of word sequences, showing the cortical tracking of sentence structure (Fig. 2). Such dissociation, as discussed in the following, cannot be achieved by typically contrasting sentence and word list (Burroughs et al., 2021; Hultén et al., 2019; Woolnough et al., 2023). We further uncover the temporal profiles of this processing (Fig. 3). These results suggest that low-frequency neural response subserves the encoding of word property and plays a key role in sentence parsing.

Word property is the foundation to construct sentences. It has been found that PoS information could be decoded during continuous speech (Gwilliams et al., 2023). On the other hand, recent studies using frequency tagging show no significant neural tracking of PoS (Burroughs et al., 2021; Jin et al., 2020; Lo et al., 2022). However, the current study finds the representation of PoS contributes to a portion of low frequency neural responses (Fig. 3). The potential explanation of this contradiction lies in the sentence context. In the former decoding study and canonical early left-anterior negativity (ELAN) effect (Herrmann et al., 2011), as well as the current study, words are presented with sentence context. In polysemy situations, especially, the part-of-speech can be decoded from the top-down context instead of the bottom-up part-of-speech distribution (Gwilliams et al., 2023). However, in other studies, words are presented in word lists, and the representation of words might be relatively shallow or weak without the demand of sentence comprehension (Burroughs et al., 2021; Jin et al., 2020; Lo et al., 2022). The current study also reveals the temporal dynamics of PoS representation through multivariate analysis (Fig. 3b). It is noteworthy that the experiment design enables the decorrelation between word property and sentence structure, making the analysis more reliable. The neural dissimilarity between nouns and verbs peaked at around 120 ms and 490 ms, in accordance with the time window of ELAN (between 120 ms and 200 ms) and late syntactic integration reported previously (Friederici, 2011). Our results build on and extend these violation-based studies into natural speech. Interestingly, this process is slightly earlier than the phonetic processing at around 140 ms (Fig. S2c), providing further evidence about the top-down processing of PoS. This parallel representation of low-level and high-level linguistic features is in line with recent investigations about the parallel prediction of multiple levels (Brodbeck et al., 2022).

Beyond word properties, neural tracking of sentence structure is a widely observed phenomenon that provides a mechanism to temporally integrate words into larger structures (Bonhage et al., 2017; Ding et al., 2016; Lerner et al., 2011; Norman-Haignere et al., 2022; Pallier et al., 2011). The cortical dynamics tracking the sentences slowly changed over the time course (Fig. 2a, Zhang & Ding, 2017; Zhou et al., 2016), allowing the dissociation of representation at each position (Fig. 3b). The transient neural representation of words is therefore locked and nested at different phases or amplitudes of the sentential-rate cortical dynamics. This concurrent neural tracking of sentence structure and word property might be a potential mechanism of the interaction between different language hierarchies. The prediction of word property can be generated according to the phase of sentence structure to facilitate top-down processing (Arnal & Giraud, 2012; Giraud & Poeppel, 2012).

Furthermore, low-frequency neural tracking might modulate higher-frequency neural oscillations (Brennan & Martin, 2019; Giraud & Poeppel, 2012; Murphy, 2024). It has been found that the broadband gamma oscillation progressively increased across sentences, suggesting additional representations or operations underpinning sentence structure building (Desbordes et al., 2023; Fedorenko et al., 2016; Hultén et al., 2019; Woolnough et al., 2023). These representations or operations might also recruit slow cortical dynamics (Pylkkänen, 2019). In this study, the representation of word order also has a peak at around 300∼400 ms, which can be explained by additional syntactic binding or semantic composition besides the sustained structure tracking that further drives the representational difference between words with respective order.

Other studies use word list as contrast to indicate the specific significance of structure building that word property or word relatedness cannot fully account for the sentential-rate response (Burroughs et al., 2021; Lu et al., 2023). These studies demonstrate language processing beyond single words, but sentence context might yet facilitate or recruit the processing of words (Woolnough et al., 2023). The current study contributes to this obstacle through a new method to factorize sentence comprehension, in which low-level word processing is neatly controlled. This dissociation also helps characterize the neural correlates of individual cognitive differences. The strength of the sentential-rate neural response is sensitive to the proficiency of language (Ding et al., 2016; Xu et al., 2023) and can even predict the performance of sentence comprehension (Ding et al., 2017). In this study, we find the strength of sentence structure tracking is related to the task performance, while the processing of word property might be more fundamental or ceiled. More generally, the coupling between the processing of chunks and the cues to parse these chunks is a universal challenge in cognitive neuroscience. This study provides a dissociation method that can be used in broader situations.

## Materials and methods

### Participants

Twenty native listeners of Mandarin Chinese (10 females and 10 males, mean age 21.2 ± 2.13 years) participated in the MEG experiment. All subjects were right-handed with normal hearing. Two participants were excluded in the analysis since they had behavior results lower than the chance level. In compliance with institutional guidelines, all subjects gave written informed consent before enrollment and received monetary compensation. The experimental procedures were approved by the Ethics Committee of the College of Biomedical Engineering and Instrument, Zhejiang University (No. 2022–001).

### Experiment design

#### Stimuli and task

We constructed 24 sequences in Mandarin Chinese with each sequence containing 37 disyllabic words. From the second word in each sequence, every 3 words compose a subject-verb-object sentence; while from the first word, every 3 words compose an object-subject-verb sentence (Fig. 1a). Thus, 24 sequences form 24 trials for SVO and OSV conditions with each trial containing 12 sentences. However, the sentence meanings are not paired in SVO and OSV so the structural effect might be confounded by this effect. For instance, “school organized parties” and “meetings school organized” have different components “parties” and “meetings” and might activate different neural responses (Fig. 1a). To deal with this potential obscure, we reversed the word sequences and generated reversed trials, in which we had “parties school organized” and “school organized meetings” corresponding to “school organized parties” and “meetings school organized” (Fig. S1). In total, there were 48 trials in each condition. Then, we selected 12 trials (6 original and 6 reversed) and changed the words to make them semantically abnormal to generate outlier trials. Additionally, 5 extra trials for each condition were constructed in the same way for the practice session.

We synthesized the speech of these sentence sequences as stimuli. All syllables were independently synthesized and were adjusted to the same intensity and the same duration, i.e., 250 ms (Ding et al., 2016). The synthesized syllables were directly concatenated. Therefore, words and sentences were isochronously presented at 2 Hz and 2/3 Hz respectively without any prosodic cues.

#### Experiment procedures

In the experiment, neural responses were recorded using MEG. The experiment consisted of two conditions SVO and OSV that were mixed in a randomized order. SVO and OSV presented sentence sequences with the word order of “subject-verb-object” and “object-subject-verb” respectively. Each condition contains 36 normal trials and 12 outlier trials. In a trial, participants heard a starting tone first and then heard the sentence sequence after a random interval between 1s and 2s. Participants were required to follow the corresponding word order and understand the sentence sequence. Participants closed their eyes during the listening. After listening to the speech, participants were asked to report whether there were abnormal sentences in the sequence. Only normal trials were used in MEG analysis. Participants were familiarized with the task and went through a practice session before the experiment. The practice session ended after the participants made at least 8 correct responses in 10 practice trials.

### Data Acquisition and Pre-processing

Neuromagnetic responses were recorded using a 306-sensor whole-head MEG system (Elekta-Neuromag, Helsinki, Finland) at Shenzhen University, sampled at 1 kHz. The system had 102 magnetometers and 204 planar gradiometers. Four MEG-compatible electrodes were used to record EOG at 1kHz. To remove ocular artifacts in MEG, the horizontal and vertical EOG were regressed out from the recordings using the least-squares method. Four head position indicator (HPI) coils were used to measure the head position inside MEG. The positions of three anatomical landmarks (nasion, left, and right pre-auricular points), the four HPI coils, and at least 200 points on the scalp were also digitized before the experiment. For MEG source localization purposes, structural Magnetic Resonance Imaging (MRI) data were collected from all participants using a Siemens Magnetom Prisma 3 T MRI system (Siemens Medical Solutions, Erlangen, Germany) at Shenzhen University. A 3-D magnetization prepared rapid gradient echo T1-weighted sequence was used to obtain 1×1×1 mm^3^ resolution anatomical images.

Normal and outlier sequences were mixed and presented in a random order. Only the normal trials were analyzed. Temporal Signal Space Separation (tSSS) was used to remove the external interference from MEG signals (Taulu & Hari, 2009). Since the current study only focused on responses at 2/3 Hz, 2 Hz, and 4 Hz, the MEG signals were bandpass filtered between 0.3 and 40 Hz using a linear-phase finite impulse response (FIR) filter (−6dB attenuation at the cut-off frequencies, 10 s Hamming window), and downsampled to 100 Hz sampling rate.

### MEG Source Localization

Source activity was estimated by computing linear constrained minimum variance (LCMV) beamformer coefficients for each participant using the MNE Python toolbox (Gramfort et al., 2013; Van Veen et al., 1997). Volume conduction estimates were based on individual structural T1 scans, using single-layer boundary element models (BEMs) from the FreeSurfer (Dale et al., 1999; Fischl, 2012). The source space comprised 4,098 source points per hemisphere. MEG/MRI alignment was performed based on fiducials and digitizer points along the head surface (iterative closest point procedure after initial alignment based on fiducials). The sensor covariance matrix used was computed across all trials and the lambda regularization parameter was set to 5%. The individual LCMV cortical actiavtions were morphed to the ‘fsaverage’ template brain provided by FreeSurfer.

### Data analysis

#### Frequency-domain Analysis

In frequency-domain analysis, to avoid the influence of onset response, the responses during the first sentence were removed for every trial (Lu et al., 2022). To ensure that sentences are the same in SVO and OSV (Fig. S1), we also removed the last sentence in each trial. For SENTENCE, we calculated the difference between SVO and OSV but using OSV shifted one word (i.e., 500 ms) ahead. In this setting, the word properties in SVO and OSV are strictly aligned, including lexical semantics, phonological features, PoS (e.g., noun or verb), and argument roles (e.g., subject or object). Since the previous words are the same, we control the semantic dissimilarity (Broderick et al., 2018; Lu et al., 2023). Moreover, we used GPT-2 (Radford et al., n.d.) to compute the lexical predictability of each word, which is characterized by surprisal (i.e., the negative log of the probability of each word based on the context), and found the lexical predictability is also controlled (Fig. 1b). Consequently, the neural response was 15 s in duration for each trial and each condition.

The average of all trials was transformed into the frequency domain using the Discrete Fourier Transform (DFT) without any additional smoothing window. The frequency resolution of the DFT analysis was 1/15 Hz. In sensor-level analysis, the DFT was separately applied to each gradiometer sensor. For the visualization of topography, the responses of the gradiometers at the same position were averaged after the DFT. In source-level analysis, the DFT was separately applied to each vertex.

We normalized the response amplitude at 2/3 Hz and 2 Hz by subtracting the 1/f spectrum noise of spontaneous neural activity. The spectrum noises were estimated by the average response amplitude of 4 neighboring frequency bins (Ding et al., 2017). In sensor-level analysis, the statistical significance of a spectral peak averaged over sensors was tested through a one-sample *t*-test for the normalized amplitude. The significance of a spectral peak at each sensor was tested using a permutation t-test across 5000 permutations. The significance test was applied to the response amplitude at the 2/3 Hz and 2 Hz. False discovery rate (FDR) correction was applied in the two tests. In lateralization analysis, we calculate the lateralization index as LI = R / (L+R), in which L and R refer to the normalized 2/3 Hz responses of the left and right sensors. Responses of individual sensors smaller than 0 are rectified to 0. The significance of lateralization was tested using a one-sample *t*-test with FDR correction. In source-level analysis, the significance of a spectral peak was tested using a cluster-based permutation test, with cluster forming using threshold-free cluster enhancement (TFCE) and the number of permutations = 10000 (Smith & Nichols, 2009). In correlation analysis, we calculated the Spearman correlation between sensor-level response amplitude and task recall (the number of true outlier trials divided by the number of all outlier trials). The significance of correlation analysis was corrected using FDR correction.

#### Representational Similarity Analysis (RSA)

The neural representational dissimilarity matrices (RDMs) were calculated for each participant (Fig. 3a). Each entry in the neural RDMs represents the correlation distance (i.e., 1 – Pearson r) between the neural responses of two words (Walther et al., 2016). Before the computation of neural RDMs, the neural responses were averaged over trials. For each condition, we got two averaged neural responses (one averaged over origin trials and one averaged over reversed trials). For each averaged neural response, the first and the last sentences were removed with 10 sentences left. In total, we had 3×10×4 words. We computed the pair-wise correlation distance, which resulted in 120×120 neural RDMs. The 120 words can be divided into 6 categories, S_1_, V_2_, O_3_, O_1_, S_2_, and V_3_, which is referred to by its argument structure and its order in the sentence. As the word duration is 500 ms and the neural responses were sampled at 100 Hz, we had 50 time points. At each time point, we calculate a neural RDM. Only 204 gradiometers were used.

Three model RDMs were proposed to test the representation of neural RDMs, termed the order model, the part-of-speech model, and the argument model (Fig. 3a, right). The order model tests the representation of sentence structure that words at different positions have different neural responses. For word property, two kinds of features will generate sentential-rate responses. The part-of-speech model tests the representation that words with different lexical categories have different neural responses. The surprisal model tests the representation that words with different level of predictability have different neural responses (Goldstein et al., 2022). The surprisal is calculated using the GPT-2 model previously. Additionally, two control RDMs were considered (Fig. S2a). The condition model controls the representation that the words in SVO and OSV have different neural responses. The phonetics model controls the representation of the phonetic features (Mesgarani et al., 2014), which is calculated by the correlation distance of Mandarin phonetic features in the International Phonetic Alphabet (IPA).

Considering the correlation between model RDMs (Fig. S2b), the model fit was estimated using partial Pearson correlation at each time point. The significance of model fit was tested using a cluster-based permutation test. The noise ceiling was assessed using leave-one-out cross-validation, which computed the average Pearson correlation between the RDM for each subject and the average RDM over the remaining subjects (Nili et al., 2014). For the PCA visualization, we computed the first two components of RDMs averaged across participants at each time point.

## Acknowledgments

We thank the Magnetic Resonance Imaging Center of Shenzhen University for its assistance in data collection. This work was supported by the National Key Research and Development Program of China (No. 2021ZD0204105).

## Data and code availability

All data and code will be deposited at Zenodo (https://zenodo.org/doi/10.5281/zenodo.11442095) and is publicly available as of the date of publication.

## Supplemental information

Document S1. Figures S1-S2

