## Supplemental Information for "Low-frequency Cortical Activity Reflects Context-dependent Parsing of Word Sequences"

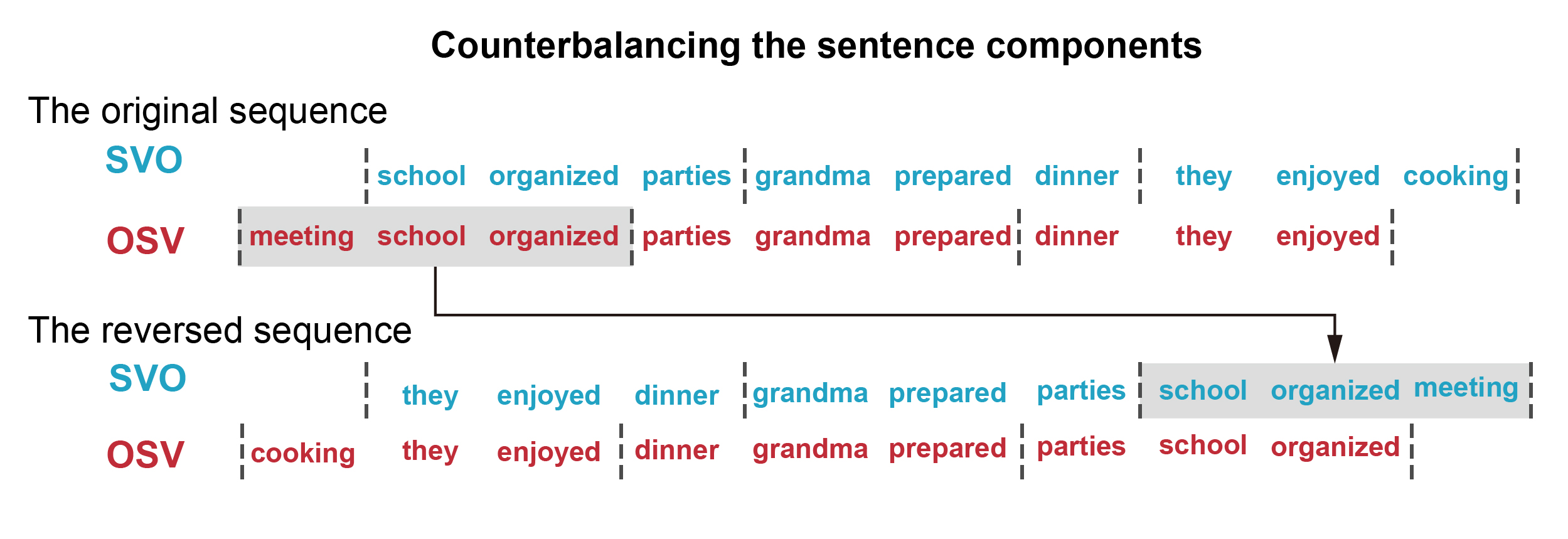
Fig. S1. Counterbalancing the sentence components. To control the sentences in SVO and OSV having the same components, we reversed the sequence but kept the local order of subject and verb in a sentence. Thus, for example, for “meeting school organized” in OSV, we also have “school organized meeting” in SVO.

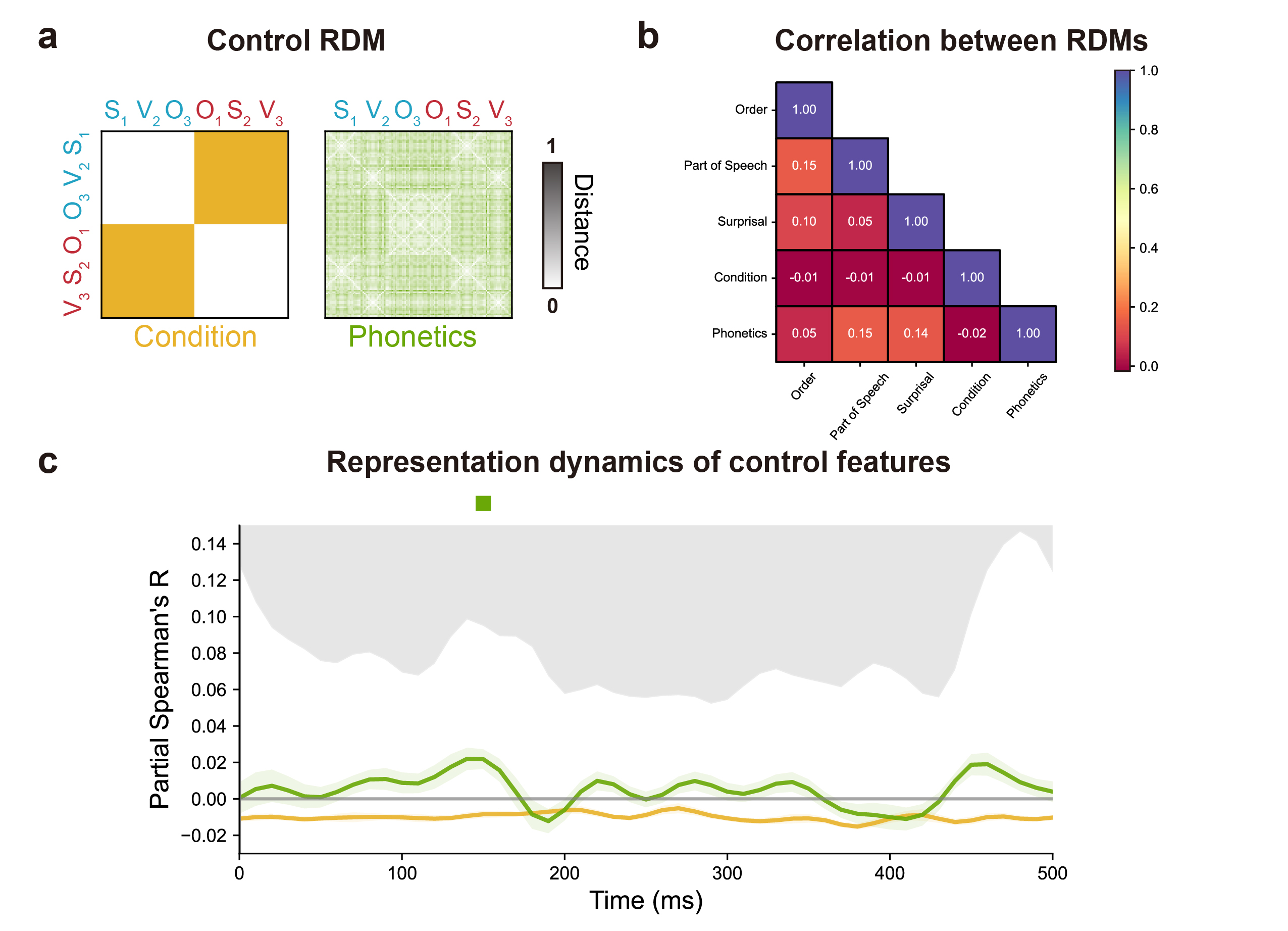
Fig. S2. **a** The control RDMs. **b** The Pearson’s correlations among model RDMs and control RDMs. **c** Time-resolved model fit for control RDMs.
